# Overcoming the widespread flaws in the annotation of vertebrate selenoprotein genes in public databases

**DOI:** 10.1101/2024.10.30.620813

**Authors:** Max Ticó, Emerson Sullivan, Roderic Guigó, Marco Mariotti

## Abstract

Selenocysteine (Sec) is a non-canonical amino acid incorporated into selenoproteins, oxidoreductase enzymes carrying essential roles in redox homeostasis. Sec insertion occurs in response to UGA, normally interpreted as stop codon, but recoded in selenoprotein mRNAs. Owing to the dual function of UGA, the identification of selenoprotein genes poses a challenge.

We show that the vertebrate selenoprotein genes are widely misannotated in major public databases. Only 12% and 6% of selenoprotein genes are well annotated in Ensembl and NCBI GenBank, respectively, due to the lack of dedicated selenoprotein annotation pipelines. In most cases (81% and 84%), overlapping flawed annotations are present which lack the Sec-encoding UGA. In contrast, NCBI RefSeq employs a dedicated selenoprotein pipeline, yet with some shortcomings: its selenoprotein annotations are correct in 76% of cases, and most errors affect families with a C-terminal Sec residue.

We argue that selenoproteins must be correctly annotated in public databases and that must occur via automated pipelines, to keep the pace with genome sequencing. To facilitate this task, we present a new version of Selenoprofiles, an homology based tool for selenoprotein prediction that produces predictions with accuracy comparable to manual curation, and can be easily deployed and integrated in existing annotation pipelines.

## INTRODUCTION

Selenoproteins are a group of proteins that contain the amino acid selenocysteine (Sec or U), the twenty-first amino acid in the genetic code. Sec is co-translationally inserted via recoding of UGA, a codon which normally terminates protein synthesis (1, 2). Sec insertion primarily depends on a *cis-* acting RNA structure called SECIS element, located in 3’ UTR regions of selenoprotein mRNAs of eukaryotes (3). The human selenoproteome consist of 25 selenoprotein genes, which have important roles in various biological processes including redox homeostasis, hormone maturation, immune response, and others (2, 4). Thioredoxin reductases (TXNRD), glutathione peroxidases (GPX) and iodothyronine deiodinases (DIO) are represented by 3, 5, and 3 selenoprotein genes in human, respectively, and they are arguably the most well studied eukaryotic selenoprotein families. TXNRDs and GPXs are essential components of the two major antioxidant systems in mammals, the thioredoxin and glutathione system respectively, while DIOs control thyroid hormone maturation. The functions of approximately a third of human selenoproteins are yet uncharacterized or poorly defined. Most selenoprotein families include homologs that substitute Sec with cysteine (Cys), i.e. “Cys homologs”. These may be found as orthologs (e.g. *Drosophila* thioredoxin reductase*Trxr1*) and/or as paralogs (e.g. human *GPX5, GPX7, GPX8*).

Selenoprotein genes have been identified in diverse branches of the tree of life. In this work, we focus on eukaryotes only. Vertebrate have relatively well-characterized selenoproteomes encompassing 19 protein families, some represented by several homologs per genome. Tetrapods typically encode 24–25 selenoproteins, whereas fish show a broader variation, with counts ranging from 28 to 55 (5, 6). Sec usage was also observed in other eukaryote species, including most metazoa, algae, some protists, and a handful of fungi (7–9). On the other hand, Sec is absent in higher plants, yeasts, many insects and few nematodes (7, 9–11).

Owing to the rare dual function of the UGA codon, selenoproteins are usually neglected by standard gene annotation programs, so that specialized software is required (7, 12). Relevantly for this work, our group previously developed Selenoprofiles, an automated pipeline to predict selenoprotein genes by homology to a set of built-in profiles representing known selenoprotein families (13, 14).

In the present work, we assessed selenoprotein gene annotation in two major public databases, Ensembl and NCBI, We focused on vertebrates, arguably the best annotated of all genomes. By using Selenoprofiles to predict selenoproteins in genome assemblies and assessing the corresponding gene annotations, we show evidence for widespread misannotation of selenoprotein genes.

To remedy this important issue, we have upgraded Selenoprofiles with several key enhancements: a streamlined, one-command installer; an automatic orthology-assignment utility that labels each prediction with gene names; and a lineage-aware filter that retains only candidates consistent with the expected selenoproteome. Together, these features yield results that match the precision of manual curation, and facilitate the integration of Selenoprofiles in automated genome annotation pipelines.

## MATERIAL AND METHODS

### Genomic sequences and resources

We downloaded all assemblies and annotations available in Ensembl version v.108 (15), comprising of 315 genomes, each corresponding to a different species or strain. For the data at the National Center for Biotechnology Information (NCBI), we used the tool “NCBI datasets” to obtain all genome assemblies corresponding to vertebrates, then considered only those with an available gene annotation (16, 17). Because some annotations were available but limited to the mitochondrial genome or scaffold structure, we only considered GTF files whose archive size was >3MB. This resulted in a set of 1,224 NCBI vertebrate genomes, each representing a distinct species. For analysis, we differentiated between RefSeq (ids starting with “GCF_”) from GenBank (“GCA_”) entries. The phylogenetic tree of Ensembl species we downloaded from Ensembl Compara (18). The analogous NCBI tree was obtained from NCBI Taxonomy (19).

### Selenoprotein gene prediction

Selenoprotein gene prediction was performed using Selenoprofiles (13, 14) version v4.5.0, available at https://github.com/marco-mariotti/selenoprofiles4. Selenoprofiles is a computational pipeline to identify members of the known selenoprotein families and related proteins. It uses several homology-based gene prediction programs, and employs a set of manually curated multiple sequence alignments of selenoprotein families to scan genomes for homologues. We used 21 profiles representing 19 protein families (some families encompass orthologous groups that are distant enough to warrant different profiles, for better prediction performance). The profiles are available at https://github.com/marco-mariotti/selenoprotein_profiles (v1.4.0). We scanned genome assemblies for all known selenoprotein families in metazoa and focused our analyses on the predictions labelled as “selenocysteine”, i.e., the genes encoding for selenoproteins, with no apparent pseudogene features (e.g. frameshifts).

### Annotation assessment

To assess the annotation of selenoproteins in public databases, we developed a new utility called *Selenoprofiles assess*, now integrated in Selenoprofiles v4.5.0. This tool takes as input two GTF/GFF files, one from Selenoprofiles and one from another source, and matches gene annotations based on their overlap in genomic coordinates. Overlapping annotations must occur on the same strand, and on the same frame (except for the “Out of frame” classification). After analysis of the Sec-encoding codon and the rest of coding sequence, each Selenoprofiles prediction is assigned an annotation status label, as shown in Figure 1 and described in Results. To make this script, we took advantage of PyRanges, a package to represent and manipulate genomic data in Python (20). Plots were built using the R package ggplot2 (21).

**Figure 1:**
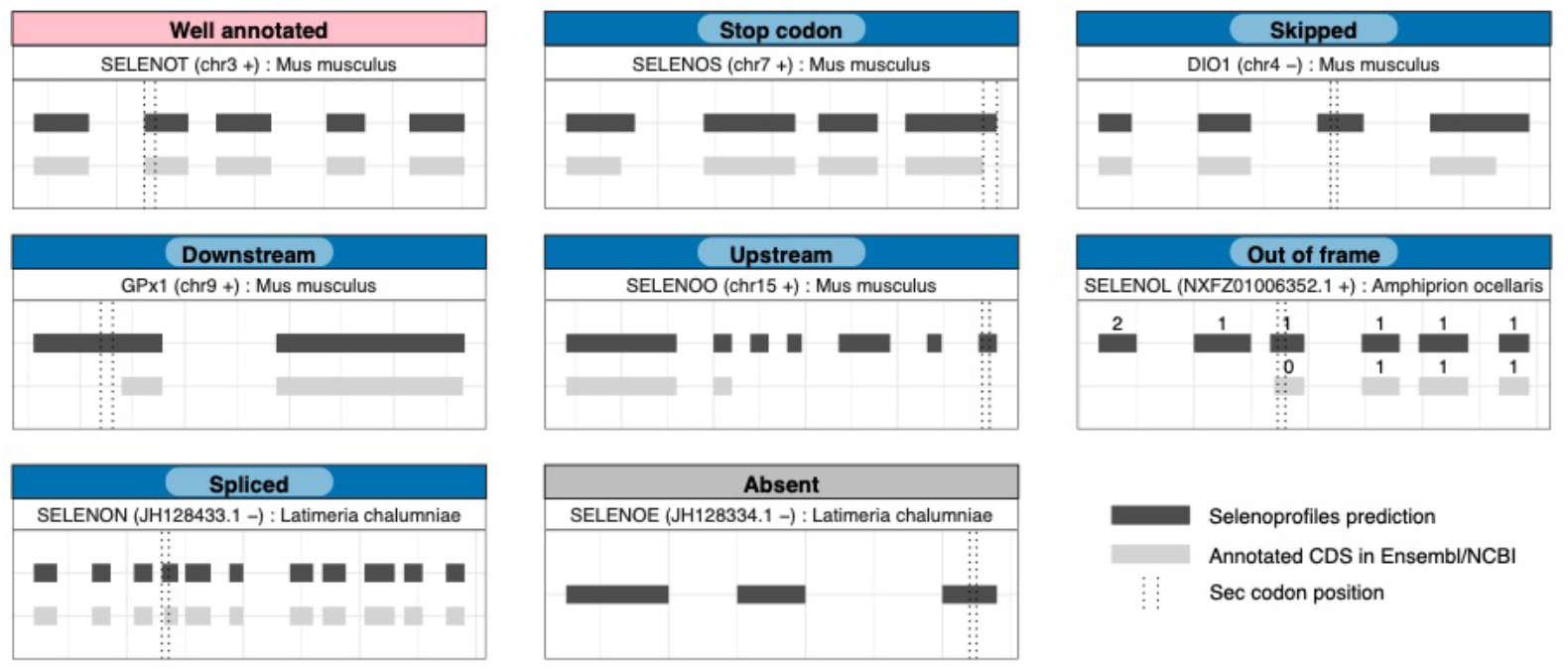
Classification of selenoprotein annotation/misannotations. Segments represent coding sequence (CDS) regions of selenoprotein genes. Selenoprofiles predicted CDS are colored in dark grey, and annotated Ensembl/NCBI transcripts are in light grey. Dotted vertical lines represent start and end positions of the Sec encoding codon. Introns were shrunk to a fixed size for visualization purposes. The classes in blue, i.e. all other than “Well annotated” and “Absent”, are collectively referred to with the term “Misannotation” in this manuscript.

### Phylogenetic reconstruction and orthology

The protein sequences of all Selenoprofiles predictions across all genomes were collected using the *Selenoprofiles join* utility. These were used as input to phylogenetic reconstruction workflows, available in the ETE3 v3.1.3 build framework (22). We ultimately present trees obtained with the “eggnog41” workflow (23), which internally uses PhyML v20160115.patched for phylogenetic reconstruction (24). We visualized gene trees with the *show_tree* program in the biotree_tools package v0.0.6 (https://github.com/marco-mariotti/biotree_tools.git), based on the ETE3 python module (22).

For every selenoprotein family with multiple paralogs in vertebrates (GPX, TXNRD, DIO, SELENOW/V), we inspected phylogenetic trees and manually partitioned them into orthologous groups, here referred to as subfamilies, by applying the species overlap method and following maximum parsimony criteria (25). To assign subfamily names, we performed similarity searches against Uniprot and consulted relevant literature (5, 26, 27).

Next, we built a new utility called *Selenoprofiles orthology*, now available in v4.5.0, constituting a fast alternative to tree building for subfamily assignment. To this aim, we first generated reference alignments (“anchors”) for each multimember selenoprotein family including representatives of each subfamily, which were assigned to orthologous groups beforehand via phylogenetic reconstruction as detailed above. Alignments were performed using Mafft v7.45331 (28). Anchor alignments were reduced to a maximum of 4, 8, or 12 representative sequences for each subfamily for each lineage, automatically selected using TrimAl (29) to preserve sequence diversity. In the *Selenoprofiles orthology* procedure, candidate sequences are aligned to reference alignments using Mafft, and Pyaln v0.1.4 (https://github.com/marco-mariotti/pyaln.git) is used for fast calculation of similarity scores of candidate sequences against the representative sequences of each subfamily. These scores are used for subfamily assignment of each candidate. Similarity score parameters are displayed in Table S1. Results of our benchmark with 3,952 Selenoprofiles predictions in Ensembl genomes are shown in Table S2, which led to choosing anchor alignments with size of 12 sequences, and this Pyaln sequence comparison method: gap positions are excluded (gaps = n), and the average weighted sequence identity “AWSI” (metrics = ‘w’) is computed with weights based on the maximum frequency of non-gap characters (weights = ‘m’). The same similarity metrics is used for ranking gene predictions in the novel filtering procedure *Selenoprofiles lineage* (see Results). Sequence and tree manipulations throughout the project were carried out using the programs *alignment_tools* and *tree_tools* from the biotree_tools package.

## RESULTS

### Evaluation of selenoprotein annotation

We set to evaluate the status of selenoprotein annotations in Ensembl and NCBI. The identification of all selenoprotein genes in a genome is not trivial, and it is likely that many selenoprotein families remain undiscovered in less studied organisms, e.g. unicellular eukaryotes (7). To elude this problem, we focused on vertebrates: their selenoproteome has been thoroughly studied (4–6, 30–35), so that, we argue, it can be reliably identified by homology to known selenoprotein families. Our dataset consisted of paired genome assemblies and genome annotations, 315 from Ensembl and 1,224 from NCBI (Methods). We subdivided NCBI genomes into two categories: GenBank (602 entries), which includes genome and annotation data submitted directly by researchers without further curation, and RefSeq (622 entries), which comprises a non-redundant selection of genomes consistently annotated using the NCBI Eukaryotic Genome Annotation Pipeline (EGAP) (36). As *bona fide* selenoprotein annotation, we ran Selenoprofiles, which yielded 8,666 and 26,210 “raw” selenoprotein predictions in Ensembl and NCBI genomes, respectively, corresponding to 19 protein families (see Methods, Data Availability). Later on, we critically evaluated and refined this method by adding a novel *Selenoprofiles lineage* utility, detailed in later sections of this manuscript, which yielded a higher-quality set of “filtered” predictions (7,612 and 24,135 in Ensembl and NCBI, respectively).

We inferred the status of selenoprotein annotation by matching genomic coordinates from Selenoprofiles predictions with annotated coding sequence (CDS) regions and evaluating whether the Sec codon was included in the annotated CDS. If it was not the case, we further classified them into different misannotation cases depending on the position of Sec and annotated CDS (Figure 1). CDS annotations that ended precisely at the Sec UGA were classified as “Stop codon”. The misannotation classes “Upstream”, “Downstream”, and “Skipped” were applied when there were portions of annotated CDS at the 5’, 3’, or both 5’ and 3’ of the Sec UGA, respectively. Some selenoprotein annotations exhibited a distinct frame compared to Selenoprofiles predictions, and were classified as “Out of frame” cases. Transcripts which contained a splicing site in the Sec codon were classified as “Spliced”. Instances where no annotation was overlapping to a Selenoprofiles selenoprotein CDS were classified as “Absent”.

The great majority of selenoprotein families contain a single Sec residue. Notable exceptions in vertebrates are SELENOP, which contains 10 Secs in human and up to 37 in fish (37), and SELENOL, which contains two nearby Sec forming a diselenide bond (35). While the first Sec residue of SelenoP is embedded in a conserved thioredoxin-like motif, the rest are located in a Sec-rich C-terminal tail, which is particularly challenging to identify in genomes (37). For simplicity, here we considered only the first Sec residue of each selenoprotein gene to define its annotation status.

### The status of selenoprotein annotation in Ensembl

Our results show that most selenoprotein genes have defective annotations in Ensembl: only ∼11% of predicted selenoproteins present at least one “Well annotated” entry (Figure 2A). Around 7% of selenoproteins have no annotation at all (“Absent”). The majority (∼81%) are “Misannotations”, i.e. there are only flawed annotations which lack the Sec-encoding UGA. We next examined annotation at transcript level, wherein a single Selenoprofiles gene prediction may match multiple annotated transcripts, particularly in well studied model organisms. This analysis allowed us to distinguish among misannotation types (Figure 1). We observed that almost 24% of transcripts skip the Sec-UGA-containing exon, or at least its relevant portion (Figure 2B). There are also numerous cases where the annotated CDS ends at the Sec codon (18.3%), or further upstream (21.9%), or starts downstream of it (18.4%). Other cases were more rare.

**Figure 2.**
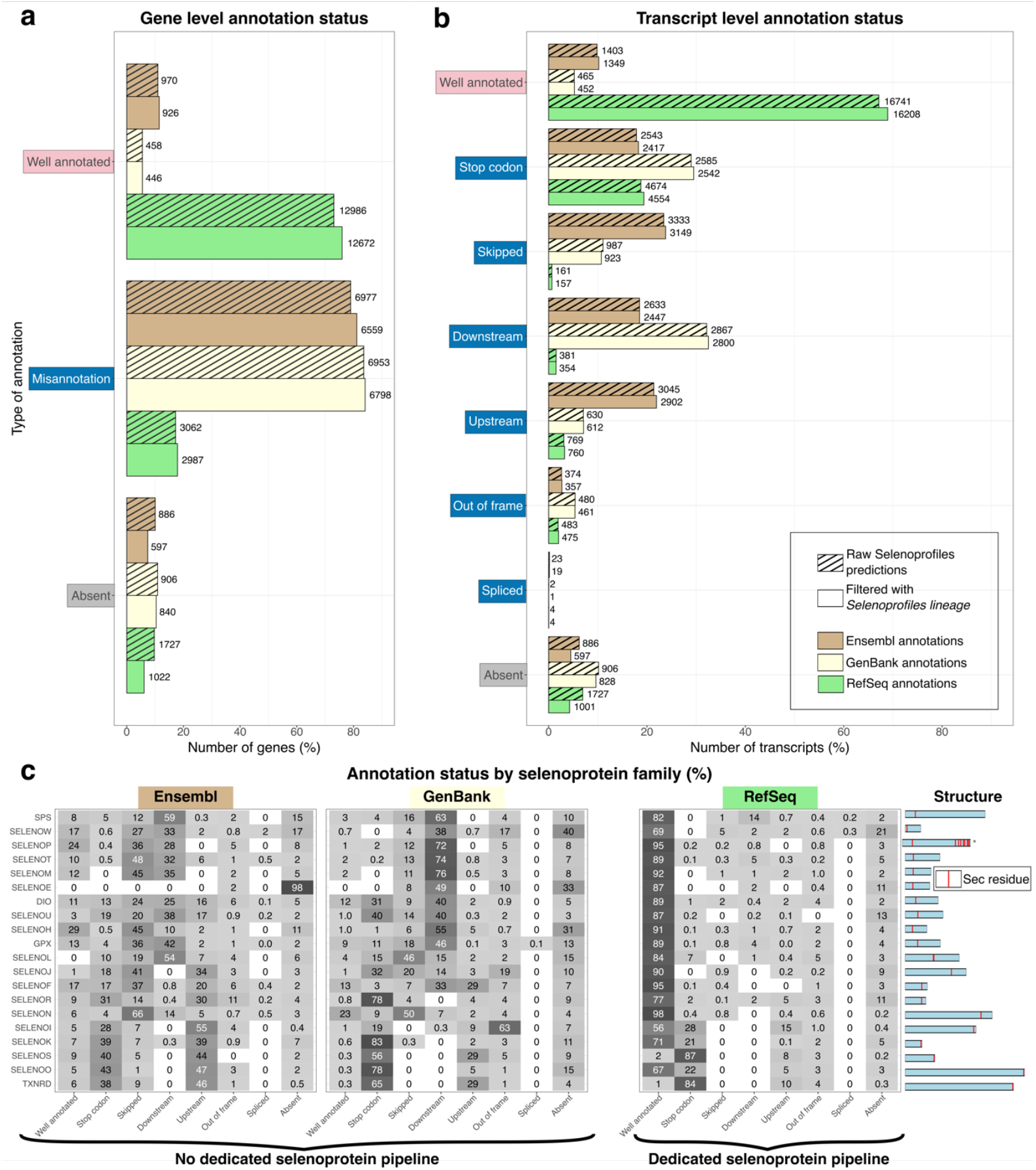
Status of selenoprotein annotation in Ensembl, RefSeq and GenBank. **a**. Bar plot showing the annotation status of selenoproteins (Selenoprofiles predictions) at gene level in different databases. Three main annotation cases are shown, with “Misannotation” aggregating all classes shown in Figure 1 other than “Well annotated” and “Absent”. Bar shading differentiates the raw (i.e. unfiltered) Selenoprofiles predictions and those filtered with Selenoprofiles lineage as explained later in the manuscript. Bar height represents percentages, and numbers on top of bars show actual counts. Bar colors correspond to each database. **b**. Bar plot showing the annotation status at transcript level in Ensembl, RefSeq and GenBank, with all possible annotation types. **c**. Heatmap representing the distribution of annotation/misannotation classes for each selenoprotein family, calculated at transcript level using filtered predictions. The right panel shows the gene structures of representatives of each family: the CDS is represented in blue, and Sec residues are displayed as red lines. Families are sorted by the position of Sec residue, with those carrying it at the C-terminus placed at the bottom. Note that the SELENOW contains both SELENOW and SELENOV genes, since they are highly homologous and included in the same Selenoprofiles profile. (*) SELENOP: Annotation status was determined using only the first Sec residue of each selenoprotein.

Next, we examined the annotation status for each selenoprotein family separately. Surprisingly, the most well-known selenoprotein families do not exhibit the best annotation status in Ensembl (Figure 2C); rather, SELENOH and SELENOP (first Sec) exhibit the highest percentage of “Well annotated” cases (29% and 24%, respectively). Consistent with intuition, we observed a clear correlation between misannotation type and the position of Sec in the gene structure. In protein families that carry Sec in close proximity to the C-terminus, such as TXNRDs and selenoproteins S, O, K, and I, the Sec residue tends to be annotated as a “Stop codon” signal, or the CDS ends further “Upstream” (Figure 2E). On the other hand, “Downstream” cases are more abundant in families containing Sec in their N-terminal part, such as GPX, DIO, SPS, and selenoproteins H, M, N, T, V, W. Unsurprisingly, the selenoproteins SELENOE (31), SELENOL (35), and SELENOJ (30), that are not present in human and mouse, exhibit the worst status in Ensembl, with <1% “Well annotated” genes.

We noticed that misannotations are not homogenously distributed across species. Instead, “Well annotated” selenoprotein genes are concentrated in few model species (e.g. human, mouse, zebrafish, pig). In contrast, almost all selenoproteins are misannotated in the rest of species. The full tree of Ensembl species with the complete classification of their selenoprotein genes is shown in Supplementary Figure S1, while Figure 3A shows a reduced version with a representative set of species. Altogether, we attribute the presence of well annotated selenoprotein genes in model organisms to the work of manual curators, while the pervasive mistakes in remaining taxa stem from the lack of a dedicated automated pipeline at Ensembl.

**Figure 3.**
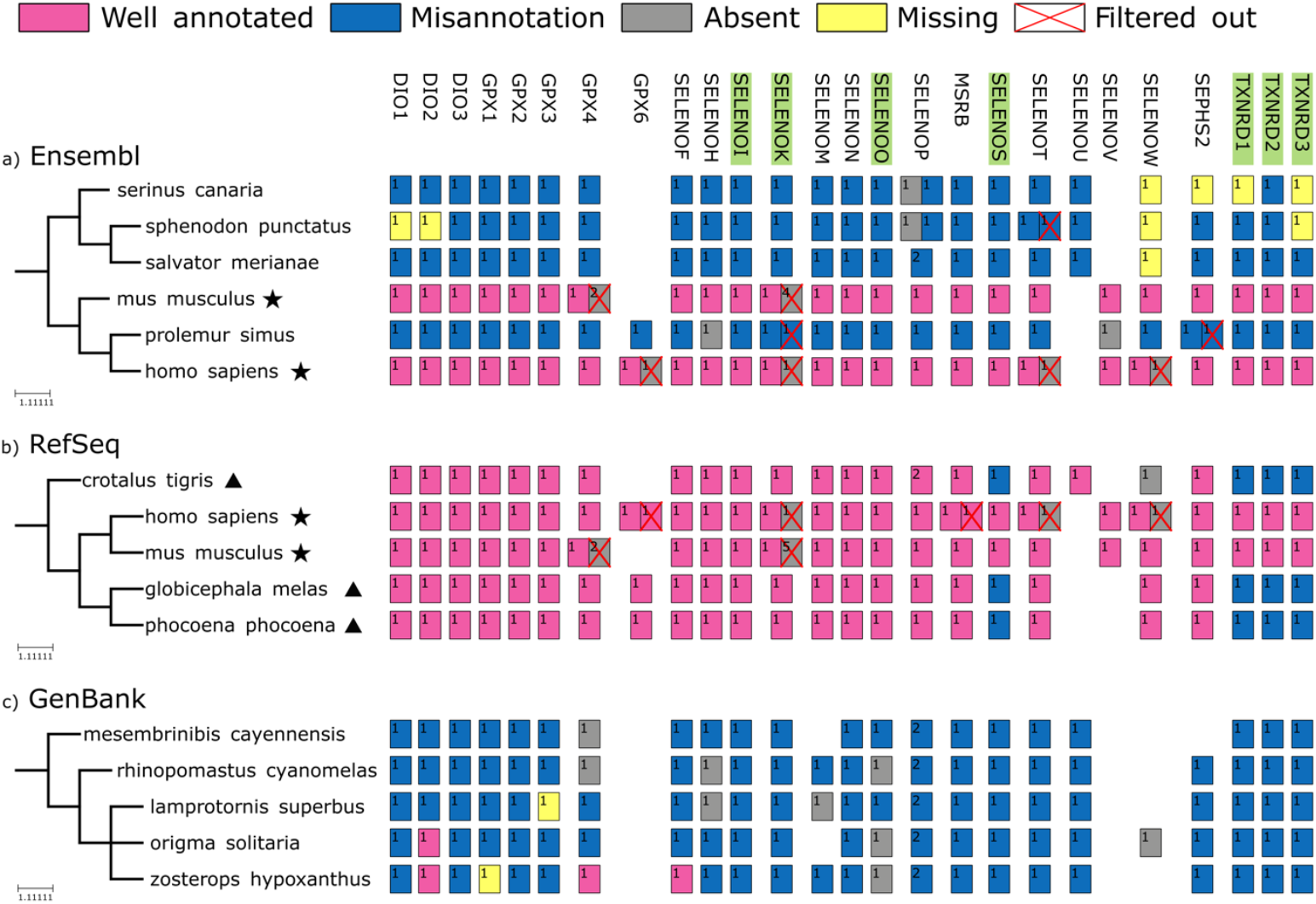
Distribution of misannotations across species. The species tree of six representative organisms is shown with the annotation of selenoprotein genes in Ensembl (**a**), RefSeq **(b)** and GenBank (**c**), color-coded by annotation/misannotation labels grouped by subfamily. Gene-level annotation classes are used (Figure 1), wherein “Absent” (grey) means that a selenoprotein predicted by Selenoprofiles has no overlapping annotation in Ensembl/NCBI. Moreover, the yellow color is used to mark “Missing” genes, i.e. expected in this genome based on lineage but not detected by Selenoprofiles. Also, red crosses mark those genes “Filtered out” by Selenoprofiles lineage, i.e. unexpected genes. Subfamilies with C-terminal Sec are highlighted in green. In the species tree, stars denote model organisms with well annotated selenoproteomes, and a triangle denote NCBI genomes with well annotated selenoproteins except for the C-terminal Sec families.

### The status of selenoprotein annotation in NCBI GenBank and RefSeq

Our analysis reveals that selenoprotein annotation is flawed in NCBI GenBank, too: only 5.5% of selenoprotein genes are “Well annotated”, while 84.1% fall into the “Misannotation” category and 10.4% are completely “Absent”. At the transcript level, the most frequent error remains treating the Sec residue as a stop codon, but the other classes are also common. Well annotated cases are spread across diverse species with no apparent underlying pattern.

Correct models appear sporadically across a wide taxonomic range with no discernible trend, reflecting the fact that GenBank records deposited by the research community are generated through diverse pipelines that do not incorporate dedicated selenoprotein routines.

In contrast, vertebrate selenoprotein genes in RefSeq genome annotations have “Well annotated” Sec residues in 76% of cases. The rest of cases are either “Absent” (6.1%) or “Misannotations” (17.9%). By analyzing the distribution of misannotation types at transcript-level and by family (Figure 2), it appears evident that the main flaw of RefSeq annotation of selenoproteins concerns the protein families with C-terminal Sec residues. Indeed, the “Stop codon” class constitutes 72.2% of “Misannotation” cases. It also appears that, in RefSeq, the annotation status of fish-specific selenoprotein families is not worse than mammalian families. Analyzing of the distribution of misannotations across species (representative tree in Figure 3; full set in Supplementary Figure S2), we notice that we can divide RefSeq genome annotations in two groups: 1. A handful of model species have selenoproteomes that are completely well annotated; 2. Some species have well annotated selenoproteins, except for a few C-terminal Sec protein families, namely SELENOS, TXNRD, and, in a lesser extent, SELENOI, SELENOK and SELENOO. Taken together, our analysis shows that RefSeq relies on manual curated entries for a few model species, while it deploys an automated pipeline for the rest. The RefSeq pipeline evidently has a procedure in-place for selenoproteins that is overall effective, but falters when the Sec residue is followed by a very short C-terminal extension.

### Improving Selenoprofiles for fully automated reliable annotation of selenoproteins

We set to provide an fully automated computational tool to produce reliable selenoprotein annotations. Previously, we developed Selenoprofiles, a pipeline for selenoprotein gene prediction in genomes featuring remarkable gene prediction accuracy (14). Yet, Selenoprofiles was not integrated in the annotation pipeline of public databases, to our knowledge. We ascribe this at least partially to the characteristics of the early versions of Selenoprofiles, which were hard to install and run. Here, we present a new version which remedied these shortcomings and introduce various other improvements. First, Selenoprofiles can now be installed via a single conda command, or by pulling a ready-made Docker image, as explained in the new comprehensive online documentation (https://selenoprofiles4.readthedocs.io/).

### *Selenoprofiles orthology*: automatic assignment of selenoprotein subfamilies

Second, we created a new utility called *Selenoprofiles orthology* to assign gene predictions to specific subfamilies, i.e. orthologous groups / paralogs. Indeed, Selenoprofiles predicts all paralogous genes of a given family (e.g. DIO1, DIO2, DIO3) using the same profile (DIO), so that, in previous versions, the user could not differentiate among paralogs without a dedicated posterior analysis. The *Selenoprofiles orthology* utility addresses this task by adopting a fast alternative to phylogenetic tree reconstruction, the state-of-the-art for orthology assignment (Figure 4). Our procedure is based on curated reference alignments referred to as “anchors”, containing selected representatives for all subfamilies for each multimember selenoprotein family. New selenoprotein predictions are aligned to anchors, and their protein sequence similarity is computed against each subfamily, ultimately assigning each prediction to its most similar subfamily.

**Figure 4.**
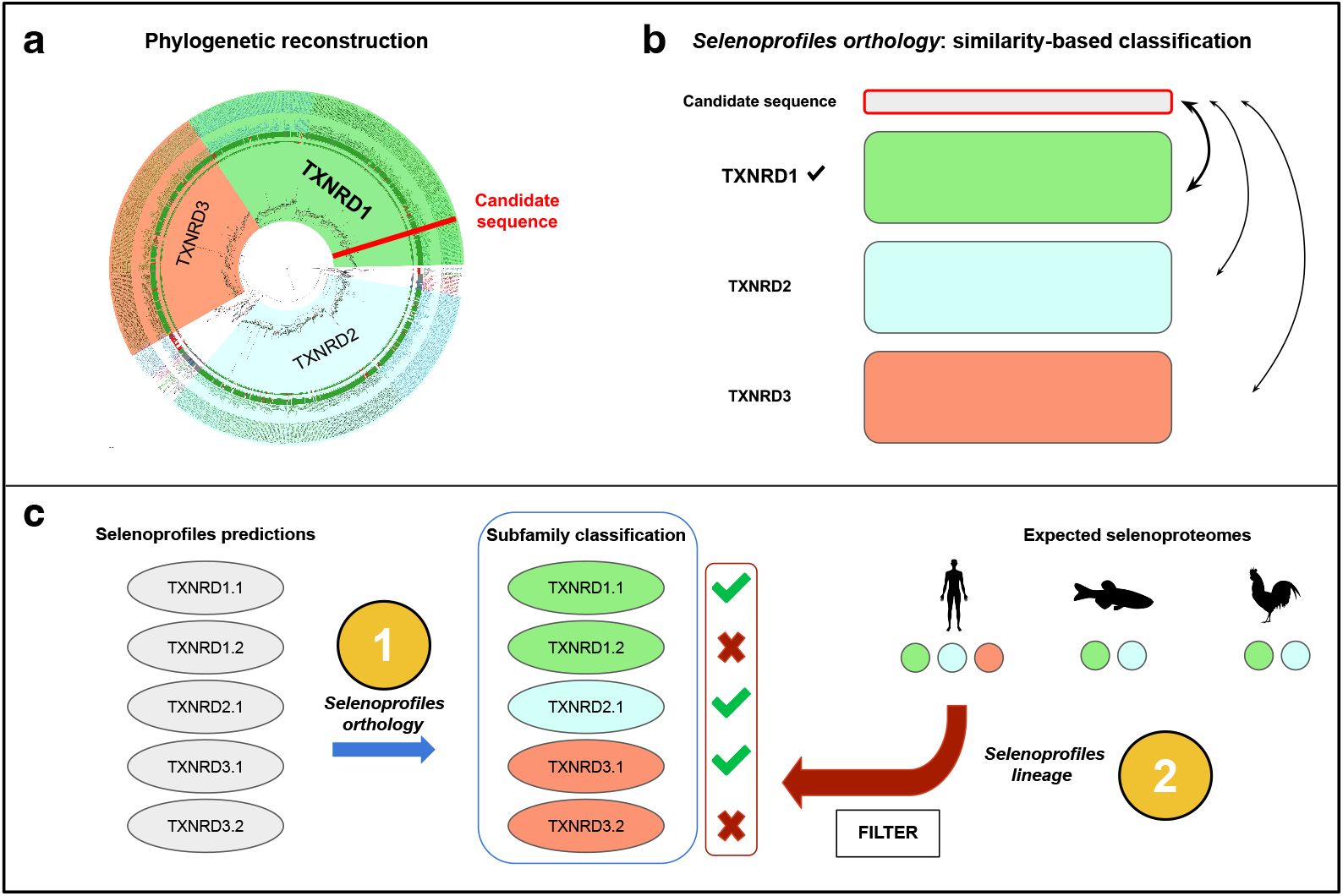
New Selenoprofiles utilities for subfamily classification pipeline and prediction refinement. Initially, we built a phylogenetic tree for every multimember family to produce a bona-fide classification of orthologous groups (**a**). Next, we devised an automated, fast similarity-based procedure that does not require tree building, now implemented as the Selenoprofiles orthology utility (**b**). Finally, we developed a filter based on the expected selenoproteome for each lineage, now implemented as the Selenoprofiles lineage utility (**c**).

To develop this functionality, we built gene trees and manually partitioned them into orthologous groups (Methods), which replicated well our previous phylogenetic analysis of vertebrate selenoproteins (5). From these, we derived anchor alignments with reliable subfamily labels, which we then reduced for faster processing by automatic selection of sequence representatives. For subfamily assignment of candidates, we tested various metrics of sequence similarity, differing by how gaps are scored and how different columns are aggregated. We then selected the method that maximized the accuracy of orthology assignment in a set of 3,952 selenoprotein predictions from four multimember families (GPX, TXNRD, DIO and SELENOW/V) in Ensembl genomes (Methods; Supplementary Tables S1 and S2).

### *Selenoprofiles lineage*: filtering based on expected selenoproteomes per lineage

Lastly, we decided to improve Selenoprofiles so that its predictions roughly match the accuracy of experienced manual curation. While our pipeline already features high sensitivity, it can lead to some false positives, typically in presence of abundant retrotransposed pseudogenes in mammalian genomes. In Selenoprofiles, most pseudogenes are already labelled and discarded due to the presence of in-frame stop codons or frameshifts (e.g. 888 predictions across 315 Ensembl genomes, an average of 2.82 per genome). Yet, some happen to lack these features and are output. At manual inspection, these may be identified by three criteria: 1. they are typically intronless, 2. most lack evidence of transcription, and 3. they are not conserved. Because some functional genes are also intronless (e.g. Sephs2 (5, 38)), the first criterion is not fit for inclusion in an automated pipeline. The second criterion is also not suitable for Selenoprofiles, because it requires additional data (transcriptomics) not universally available. We thus decided to implement the conservation criteria in a novel utility called *Selenoprofiles lineage*, which consists in a filter based on the set of selenoproteins previously observed in other species related to the one under analysis. This functionality depend on a manually curated table defining the precise selenoprotein content of all major vertebrate lineages (Data Availability). We compiled this data based on our recent analysis of vertebrate selenoproteomes (6).

*Selenoprofiles lineage* (Figure 4C) is built on top of the *orthology* utility: gene expectations are defined at the subfamily level, for improved precision. First, the species under analysis is mapped into a vertebrate lineage present in the expectation table (e.g. placentals, birds), either manually or automatically via NCBI taxonomy. Next, the program checks for extra predicted genes: for example, two copies of GPX4 are found in the mouse genome, but a single one is expected in placentals (Figure 3). Finally, genes are selected based on sequence similarity to the subfamily anchor alignment: only the N top-scoring candidates are kept, where N is the number of expected genes. Besides marking some genes as “Filtered out”, *Selenoprofiles lineage* may also report some subfamilies as “Missing”, if a given genome contains fewer predictions than expected.

Our new utility allow to refine predicted selenoproteomes in full automation. Results are shown in Figure 2, which shows the annotation status of Ensembl and NCBI before and after lineage-based filtering, and in Figure 3, and Supplementary Figures S1 and S2, wherein filtered and missing genes are shown for each species. Mostly, filtered out predictions corresponded to “Absent” and “Misannotation” rather than “Well annotated” cases, consistent with the notion that functional genes are carefully annotated more often than pseudogenes. Altogether, the *lineage* utility yields predictions with quality comparable to manual analysis. On the other hand, it may miss out on lineage-specific duplications that are not reported in the expectation table, although this was compiled via a thorough analysis of hundreds of vertebrates (6).

The new *orthology* and *lineage* functionalities are available in Selenoprofiles v4.5.0. In the future, we may expand their use to non-vertebrate genomes.

## DISCUSSION

Despite the massive advances in sequencing, genome annotation remains challenging. Due to the recoding of the UGA codon, this problem is aggravated in the case of selenoprotein genes, leading to mispredictions and omissions in public databases. The current study confirmed clear support to this hypothesis: we report that the extent of flawed annotations for selenoprotein genes reach 88% for Ensembl, 24% for RefSeq and 94% for GenBank. In most cases, an overlapping gene annotation is present but does not include the Sec codon, with many possible gene structure arrangements observed (Figures 1 and 2). In summary, accurate annotation of selenoprotein genes in Ensembl is currently largely restricted to a few model organisms; the RefSeq EGAP pipeline performs well overall, owing to their dedicated efforts to improve selenoprotein annotation (12, 36), though it consistently struggles with families bearing a C-terminal Sec residue; whereas GenBank displays widespread annotation errors across the vast majority of species and selenoprotein families.

Genome annotations are essential for understanding the information encoded in genomes. This includes elucidating evolutionary relationships, molecular functions, gene expression and regulation, among others, and non-model organisms in particular are subject of unprecedented scientific attention. Indeed, we entered an era of massive sequencing and biodiversity genomics, and in the near future the goal is to sequence all eukaryotic life on Earth, driven by initiatives under the umbrella of the Earth Biogenome Project (39–41). Naturally, manual curation cannot keep the pace of modern genomics, and having accurate automated pipelines is compelling.

It is now more important than ever that genome annotations of all organisms are as complete as possible, to enable the effective application of methodologies that make use of annotations as a key source of information. This includes a wide array of omics technologies, phylogenetic studies, and also unbiased experimental and computational approaches such as CRISPR-based screening for genes involved in a pathway of interest, and genome scans for positive selection. In these contexts, omissions and misannotations may lead to biases, incomplete design, and misguided interpretation of results.

One may think that Sec residues represent only a tiny fraction of our genomes, and may unwisely dismiss their importance. Yet, selenoprotein genes are responsible for many essential functions: most notably, they are core components of the antioxidant and redox homeostasis systems of all vertebrates, which are fundamental for life. In these enzymes, Sec is located almost invariantly in the catalytic site, where it provides a selective advantage over its canonical analog cysteine, which may be related to increased resistance to oxidative inactivation, or enhanced catalysis (2). Therefore, Sec is arguably the most important residue of many essential proteins, and its correct annotation is compelling.

In previous attempts to address the issue of selenoprotein prediction, a dedicated database was created, called SelenoDB (42, 43), now hosted at http://selenodb.crg.eu/. Yet, we reckon that a major flaw of this approach is that, in practice, only experts of selenium biology or translational recoding typically use this database. This leaves the great majority of the scientific community, which overwhelmingly uses public annotation databases, unaware and ill-equipped towards this issue. We argue that the only possible effective strategy is to correct selenoproteins in the public annotations that researchers are already using, such as Ensembl and NCBI.

To facilitate this task, we developed a new version of Selenoprofiles, a pipeline that we already showed as effective for homology-based prediction of selenoprotein in genomes (14). Selenoprofiles is now straightforward to install and run, it has a new extensive online documentation, and features new functionalities: assignment of subfamilies (i.e. precise gene naming) and lineage-based filtering, which improved specificity to a level comparable to manual curation. These are implemented for vertebrates, acting as straightforward test cases, and may be eventually extended to other eukaryotic lineages.

We advocate that the annotation pipelines used for public databases must account for the peculiar genetics of selenoproteins, and we argue that Selenoprofiles can be integrated in existing pipelines to remedy their shortcomings. We reckon that misannotations are mainly due to the in-frame UGA being penalized in the algorithms used, leading to alternative Sec-lacking gene models being selected (Figure 5). Therefore, we suggest that the output by Selenoprofiles should be fed to the combiner module that ultimately generate gene models in annotation pipelines, as an additional information layer besides the genomic data already used (e.g. for Ensembl, GeneWise predictions using Uniprot proteins and Exonerate mapping of available cDNAs)(44). Selenoprofiles only interrogates the genome sequence, while annotation pipelines typically also use transcriptomic information, so that Selenoprofiles predictions may be less accurate in regions far from Sec residue. We thus recommend that an optimal strategy would be to integrate in public annotation pipelines only the Sec encoding codons output by Selenoprofiles, prioritizing them as a high-score layer to build gene models (Figure 5).

**Figure 5.**
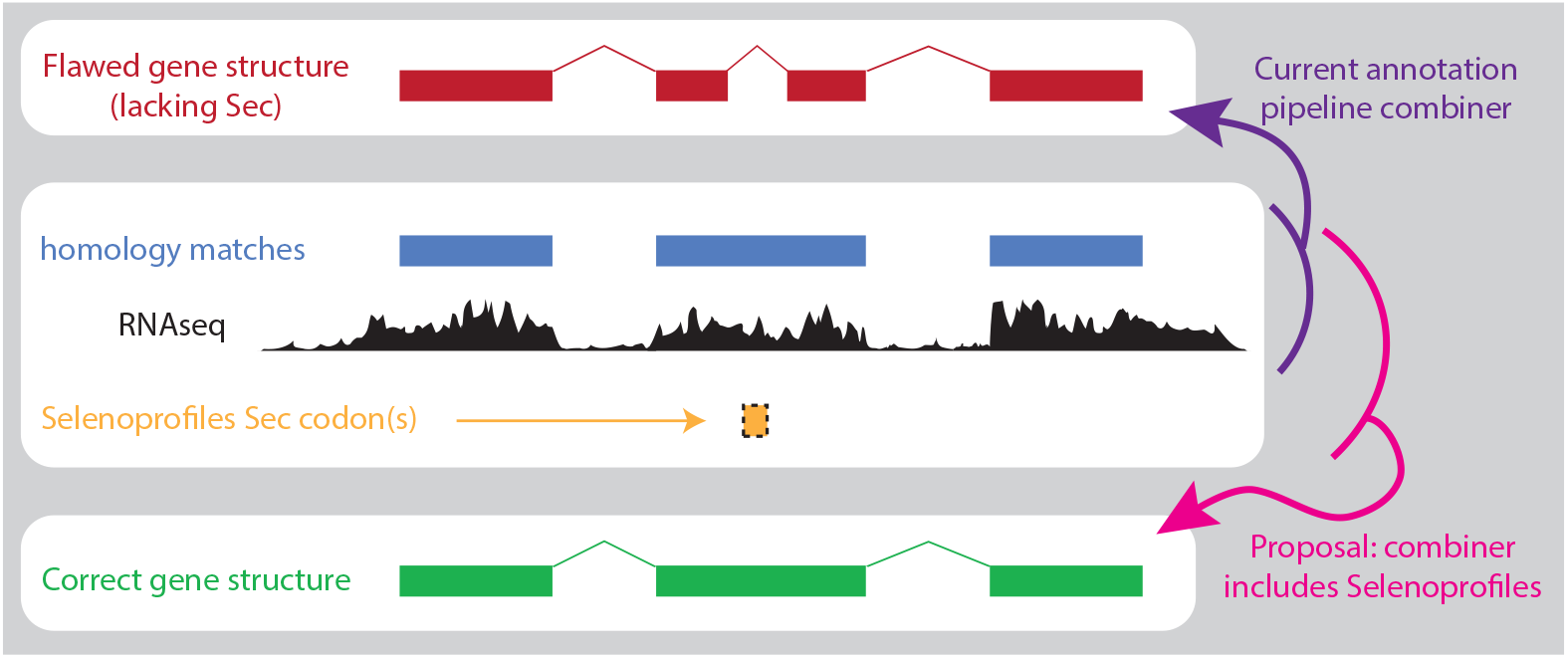
Proposed integration of Selenoprofiles in gene annotation pipelines. While existing pipelines differ in their structure, all follow the same principle in that various information layers are ultimately combined into gene structures. We propose to integrate Sec codons predicted by Selenoprofiles as high-scored layer to correct selenoprotein annotation.

Having created scalable computational tools suitable for correct annotation of selenoproteins in genomes, we will advocate for their use in public databases. We also argue that, in the near future, other well characterized cases of translational recoding (45) should receive a similar attention.

## Supporting information

All supplementary files

## DATA AVAILABILITY

The latest Selenoprofiles code is available at https://github.com/marco-mariotti/selenoprofiles4 and documented at https://selenoprofiles4.readthedocs.io/. Profile alignments, anchor alignments, and expectation tables of selenoproteins per lineage are provided as companion package https://github.com/marco-mariotti/selenoprotein_profiles. Selenoprofiles predictions on Ensembl and NCBI genomes, and related annotation status labels, can be found in Supplementary Data D1, and also online at https://mariottigenomicslab.bio.ub.edu/selenoprofiles_data/selenoprofiles_predictions/.

Supplementary Data are available online.

## SUPPLEMENTARY DATA

**Supplementary Figure S1. Selenoproteome across Ensembl genomes**. Each row represents a species and each column corresponds to a selenoprotein family. Rectangles represent individual “selenocysteine” Selenoprofiles predictions, color-labeled according to their gene annotation in Ensembl: pink for correctly annotated genes, blue for misannotated genes, grey for genes absent from the Ensembl annotation. Two graphical features are used to indicate the departure from the expected selenoproteome described in 2012 by Mariotti et al (coded in the Expectation_table.csv file at https://github.com/marcomariotti/selenoprotein_profiles v.1.2.0): yellow represent genes expected in that lineage but undetected by Selenoprofiles; and red crosses mark extra genes compared to expectations, therefore filtered by the Selenoprofiles lineage tool.

**Supplementary Figure S2. Selenoproteome across RefSeq species**. Each row represents a RefSeq species and each column corresponds to a selenoprotein family. See caption of Supplementary Figure S1 for further details.

**Supplementary Figure S3. Selenoproteome across GenBank species**. Each row represents a GenBank species and each column corresponds to a selenoprotein family. See caption of Supplementary Figure S1 for further details.

**Supplementary Table T1. Explanation of score similarity parameters optimized in Selenoprofiles orthology**. These are arguments available in the Pyaln function score_similarity. For details, see https://pyaln.readthedocs.io/en/latest/alignment.html#pyaln.Alignment.score_similarity

**Supplementary_TableT2. Results of benchmark for Selenoprofiles orthology**. See manuscript text for explanation.

## AUTHOR CONTRIBUTIONS

Max Ticó: Data curation, Formal analysis, Methodology, Visualization, Writing – original draft. Emerson Sullivan: Formal analysis. Marco Mariotti: Conceptualization, Methodology, Writing— review & editing, Supervision, Funding acquisition. Roderic Guigó: Conceptualization, Writing— review & editing.

## FUNDING

This work was supported by grants RYC2019-027746-I, PID2020-115122GA-I00, and PID2023-147164NB-I00 to MM funded by MICIU/AEI (Ministry of Science, Innovation and Universities; State Research Agency) /10.13039/501100011033 and “ESF Investing in your future”.

## CONFLICT OF INTEREST

Authors declare no conflict of interest.

## Notes

https://mariottigenomicslab.bio.ub.edu/selenoprofiles_data/selenoprofiles_predictions/

### Competing Interest Statement

The authors have declared no competing interest.

### Summary of Updates

Presubmission revision, following insightful feedback received upon our first submission to biorxiv

