## Supplementary material for "Overcoming the widespread flaws in the annotation of vertebrate selenoprotein genes in public databases": All supplementary files: SuppTables.pdf

**SUPPLEMENTARY DATA**

| | Parameters | Description |
|---|---|---|
| **Gaps** | y | Gaps considered as mismatches |
| | n | Gaps are not considered |
| | t | Terminal gaps are ignored |
| | a | Gaps are considered as any other character |
| **Metrics** | i | ASI (Average Sequence Identity) |
| | w | AWSI (Average Weighted Sequence Identity) |
| **Weights** | m | Maximum frequency of non-gap characters |
| | i | Information content |
| | q | Quadratic sum |

**Supplementary table 1. Score similarity parameters tested for *Selenoprofiles orthology*.**

These are arguments available in the Pyaln function score_*similarity*. For details, see https://pyaln.readthedocs.io/en/latest/alignment.html#pyaln.Alignment.score_similarity

| Families | Gaps | Metric | value | N_seqs | Percentage |
|---|---|---|---|---|---|
| GPx | y | ASI | 1332 | 4_seqs | 82.4767802 |
| DI | y | ASI | 1009 | 4_seqs | 100 |
| TR | y | ASI | 746 | 4_seqs | 99.2021277 |
| SelW | y | ASI | 480 | 4_seqs | 98.7654321 |
| GPx | y | ASI | 1390 | 8_seqs | 86.0681115 |
| DI | y | ASI | 1007 | 8_seqs | 99.8017839 |
| TR | y | ASI | 746 | 8_seqs | 99.2021277 |
| SelW | y | ASI | 481 | 8_seqs | 98.9711934 |
| GPx | y | ASI | 1355 | 12_seqs | 83.9009288 |
| DI | y | ASI | 1009 | 12_seqs | 100 |
| TR | y | ASI | 745 | 12_seqs | 99.0691489 |
| SelW | y | ASI | 481 | 12_seqs | 98.9711934 |
| GPx | y | AWSI.i | 1274 | 4_seqs | 78.8854489 |
| DI | y | AWSI.i | 1009 | 4_seqs | 100 |
| TR | y | AWSI.i | 737 | 4_seqs | 98.0053191 |
| SelW | y | AWSI.i | 480 | 4_seqs | 98.7654321 |
| GPx | y | AWSI.i | 1300 | 8_seqs | 80.495356 |
| DI | y | AWSI.i | 999 | 8_seqs | 99.0089197 |

| | | | | | |
|---|---|---|---|---|---|
| TR | y | AWSI.i | 745 | 8_seqs | 99.0691489 |
| SelW | y | AWSI.i | 482 | 8_seqs | 99.1769547 |
| GPx | y | AWSI.i | 1304 | 12_seqs | 80.7430341 |
| DI | y | AWSI.i | 1009 | 12_seqs | 100 |
| TR | y | AWSI.i | 745 | 12_seqs | 99.0691489 |
| SelW | y | AWSI.i | 481 | 12_seqs | 98.9711934 |
| GPx | y | AWSI.q | 1298 | 4_seqs | 80.371517 |
| DI | y | AWSI.q | 1006 | 4_seqs | 99.7026759 |
| TR | y | AWSI.q | 744 | 4_seqs | 98.9361702 |
| SelW | y | AWSI.q | 480 | 4_seqs | 98.7654321 |
| GPx | y | AWSI.q | 1309 | 8_seqs | 81.0526316 |
| DI | y | AWSI.q | 1002 | 8_seqs | 99.3062438 |
| TR | y | AWSI.q | 746 | 8_seqs | 99.2021277 |
| SelW | y | AWSI.q | 481 | 8_seqs | 98.9711934 |
| GPx | y | AWSI.q | 1322 | 12_seqs | 81.8575851 |
| DI | y | AWSI.q | 1006 | 12_seqs | 99.7026759 |
| TR | y | AWSI.q | 744 | 12_seqs | 98.9361702 |
| SelW | y | AWSI.q | 441 | 12_seqs | 90.7407407 |
| GPx | y | AWSI.m | 1318 | 4_seqs | 81.6099071 |
| DI | y | AWSI.m | 1007 | 4_seqs | 99.8017839 |
| TR | y | AWSI.m | 745 | 4_seqs | 99.0691489 |
| SelW | y | AWSI.m | 439 | 4_seqs | 90.3292181 |
| GPx | y | AWSI.m | 1320 | 8_seqs | 81.7337461 |
| DI | y | AWSI.m | 1007 | 8_seqs | 99.8017839 |
| TR | y | AWSI.m | 745 | 8_seqs | 99.0691489 |
| SelW | y | AWSI.m | 441 | 8_seqs | 90.7407407 |
| GPx | y | AWSI.m | 1330 | 12_seqs | 82.3529412 |
| DI | y | AWSI.m | 1006 | 12_seqs | 99.7026759 |
| TR | y | AWSI.m | 745 | 12_seqs | 99.0691489 |
| SelW | y | AWSI.m | 440 | 12_seqs | 90.5349794 |
| GPx | n | ASI | 1589 | 4_seqs | 98.3900929 |
| DI | n | ASI | 1009 | 4_seqs | 100 |
| TR | n | ASI | 752 | 4_seqs | 100 |
| SelW | n | ASI | 484 | 4_seqs | 99.5884774 |
| GPx | n | ASI | 1504 | 8_seqs | 93.126935 |
| DI | n | ASI | 1007 | 8_seqs | 99.8017839 |
| TR | n | ASI | 752 | 8_seqs | 100 |
| SelW | n | ASI | 484 | 8_seqs | 99.5884774 |
| GPx | n | ASI | 1591 | 12_seqs | 98.5139319 |

| | | | | | |
|---|---|---|---|---|---|
| DI | n | ASI | 1009 | 12_seqs | 100 |
| TR | n | ASI | 752 | 12_seqs | 100 |
| SelW | n | ASI | 485 | 12_seqs | 99.7942387 |
| GPx | n | AWSI.i | 1531 | 4_seqs | 94.7987616 |
| DI | n | AWSI.i | 1009 | 4_seqs | 100 |
| TR | n | AWSI.i | 751 | 4_seqs | 99.8670213 |
| SelW | n | AWSI.i | 476 | 4_seqs | 97.9423868 |
| GPx | n | AWSI.i | 1403 | 8_seqs | 86.873065 |
| DI | n | AWSI.i | 1008 | 8_seqs | 99.900892 |
| TR | n | AWSI.i | 752 | 8_seqs | 100 |
| SelW | n | AWSI.i | 480 | 8_seqs | 98.7654321 |
| GPx | n | AWSI.i | 1493 | 12_seqs | 92.4458204 |
| DI | n | AWSI.i | 1009 | 12_seqs | 100 |
| TR | n | AWSI.i | 751 | 12_seqs | 99.8670213 |
| SelW | n | AWSI.i | 481 | 12_seqs | 98.9711934 |
| GPx | n | AWSI.q | 1530 | 4_seqs | 94.7368421 |
| DI | n | AWSI.q | 1009 | 4_seqs | 100 |
| TR | n | AWSI.q | 752 | 4_seqs | 100 |
| SelW | n | AWSI.q | 485 | 4_seqs | 99.7942387 |
| GPx | n | AWSI.q | 1534 | 8_seqs | 94.9845201 |
| DI | n | AWSI.q | 992 | 8_seqs | 98.3151635 |
| TR | n | AWSI.q | 752 | 8_seqs | 100 |
| SelW | n | AWSI.q | 485 | 8_seqs | 99.7942387 |
| GPx | n | AWSI.q | 1585 | 12_seqs | 98.1424149 |
| DI | n | AWSI.q | 1009 | 12_seqs | 100 |
| TR | n | AWSI.q | 752 | 12_seqs | 100 |
| SelW | n | AWSI.q | 484 | 12_seqs | 99.5884774 |
| GPx | n | AWSI.m | 1536 | 4_seqs | 95.1083591 |
| DI | n | AWSI.m | 1009 | 4_seqs | 100 |
| TR | n | AWSI.m | 752 | 4_seqs | 100 |
| SelW | n | AWSI.m | 484 | 4_seqs | 99.5884774 |
| GPx | n | AWSI.m | 1557 | 8_seqs | 96.4086687 |
| DI | n | AWSI.m | 1007 | 8_seqs | 99.8017839 |
| TR | n | AWSI.m | 752 | 8_seqs | 100 |
| SelW | n | AWSI.m | 485 | 8_seqs | 99.7942387 |
| GPx | n | AWSI.m | 1598 | 12_seqs | 98.9473684 |
| DI | n | AWSI.m | 1009 | 12_seqs | 100 |
| TR | n | AWSI.m | 752 | 12_seqs | 100 |
| SelW | n | AWSI.m | 484 | 12_seqs | 99.5884774 |

| | | | | | |
|---|---|---|---|---|---|
| GPx | t | ASI | 1355 | 4_seqs | 83.9009288 |
| DI | t | ASI | 1009 | 4_seqs | 100 |
| TR | t | ASI | 738 | 4_seqs | 98.1382979 |
| SelW | t | ASI | 484 | 4_seqs | 99.5884774 |
| GPx | t | ASI | 1389 | 8_seqs | 86.006192 |
| DI | t | ASI | 1007 | 8_seqs | 99.8017839 |
| TR | t | ASI | 735 | 8_seqs | 97.7393617 |
| SelW | t | ASI | 484 | 8_seqs | 99.5884774 |
| GPx | t | ASI | 1431 | 12_seqs | 88.6068111 |
| DI | t | ASI | 1009 | 12_seqs | 100 |
| TR | t | ASI | 751 | 12_seqs | 99.8670213 |
| SelW | t | ASI | 485 | 12_seqs | 99.7942387 |
| GPx | t | AWSI.i | 1299 | 4_seqs | 80.4334365 |
| DI | t | AWSI.i | 1006 | 4_seqs | 99.7026759 |
| TR | t | AWSI.i | 735 | 4_seqs | 97.7393617 |
| SelW | t | AWSI.i | 484 | 4_seqs | 99.5884774 |
| GPx | t | AWSI.i | 1310 | 8_seqs | 81.1145511 |
| DI | t | AWSI.i | 1006 | 8_seqs | 99.7026759 |
| TR | t | AWSI.i | 735 | 8_seqs | 97.7393617 |
| SelW | t | AWSI.i | 484 | 8_seqs | 99.5884774 |
| GPx | t | AWSI.i | 1379 | 12_seqs | 85.3869969 |
| DI | t | AWSI.i | 1006 | 12_seqs | 99.7026759 |
| TR | t | AWSI.i | 750 | 12_seqs | 99.7340426 |
| SelW | t | AWSI.i | 485 | 12_seqs | 99.7942387 |
| GPx | t | AWSI.q | 1530 | 4_seqs | 94.7368421 |
| DI | t | AWSI.q | 1009 | 4_seqs | 100 |
| TR | t | AWSI.q | 752 | 4_seqs | 100 |
| SelW | t | AWSI.q | 485 | 4_seqs | 99.7942387 |
| GPx | t | AWSI.q | 1516 | 8_seqs | 93.869969 |
| DI | t | AWSI.q | 1002 | 8_seqs | 99.3062438 |
| TR | t | AWSI.q | 752 | 8_seqs | 100 |
| SelW | t | AWSI.q | 485 | 8_seqs | 99.7942387 |
| GPx | t | AWSI.q | 1585 | 12_seqs | 98.1424149 |
| DI | t | AWSI.q | 1009 | 12_seqs | 100 |
| TR | t | AWSI.q | 752 | 12_seqs | 100 |
| SelW | t | AWSI.q | 481 | 12_seqs | 98.9711934 |
| GPx | t | AWSI.m | 1513 | 4_seqs | 93.6842105 |
| DI | t | AWSI.m | 1009 | 4_seqs | 100 |
| TR | t | AWSI.m | 752 | 4_seqs | 100 |

| | | | | | |
|---|---|---|---|---|---|
| SelW | t | AWSI.m | 454 | 4_seqs | 93.4156379 |
| GPx | t | AWSI.m | 1401 | 8_seqs | 86.749226 |
| DI | t | AWSI.m | 1005 | 8_seqs | 99.6035679 |
| TR | t | AWSI.m | 752 | 8_seqs | 100 |
| SelW | t | AWSI.m | 481 | 8_seqs | 98.9711934 |
| GPx | t | AWSI.m | 1592 | 12_seqs | 98.5758514 |
| DI | t | AWSI.m | 1009 | 12_seqs | 100 |
| TR | t | AWSI.m | 752 | 12_seqs | 100 |
| SelW | t | AWSI.m | 481 | 12_seqs | 98.9711934 |

**Supplementary table 2. Results of benchmark for *Selenoprofiles orthology*.** See manuscript text for explanation.
