## Supplementary figures and images for "Overcoming the widespread flaws in the annotation of vertebrate selenoprotein genes in public databases"

### SuppFigureS1.species_tree_Ensembl.pdf

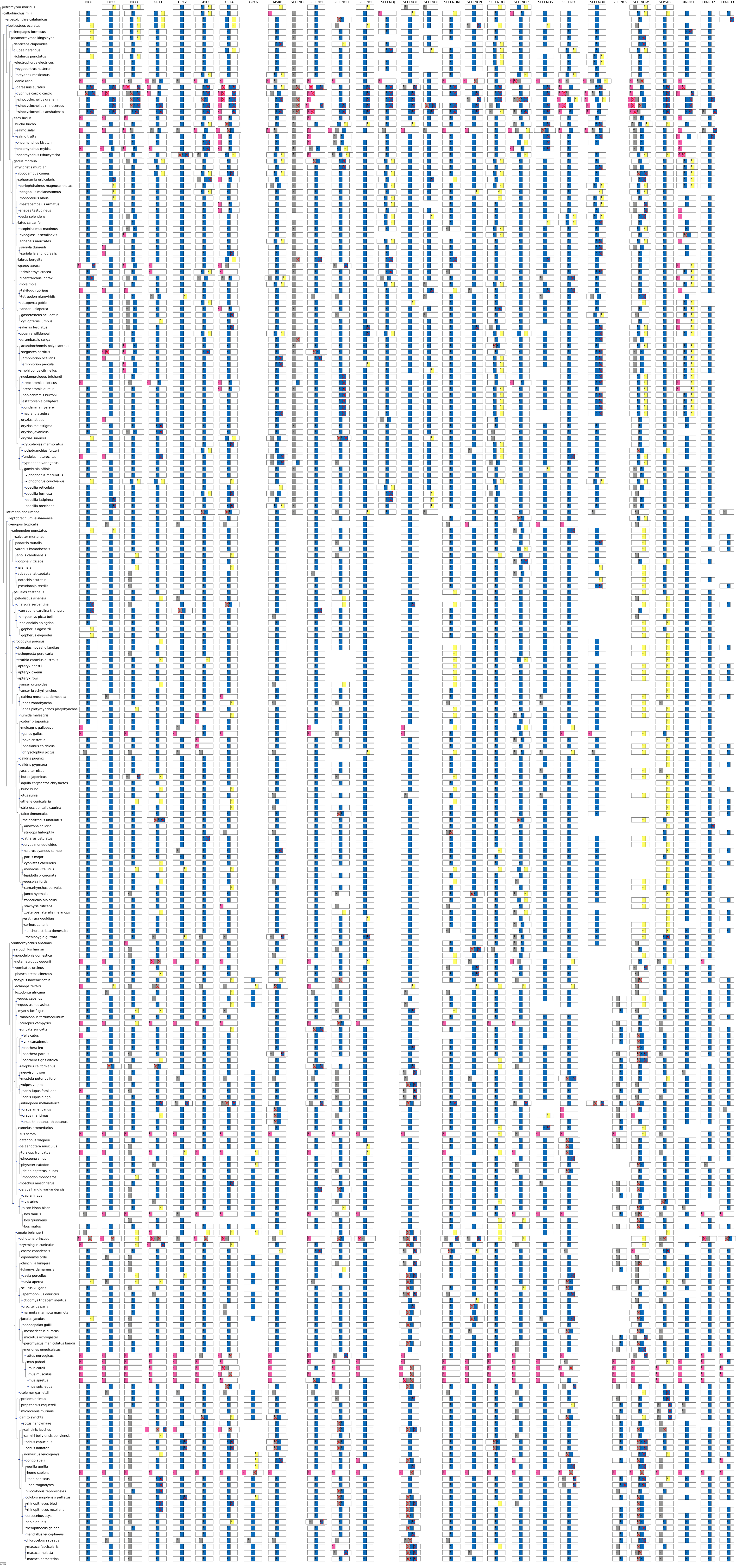

### SuppFigureS3.species_tree_GenBank.pdf

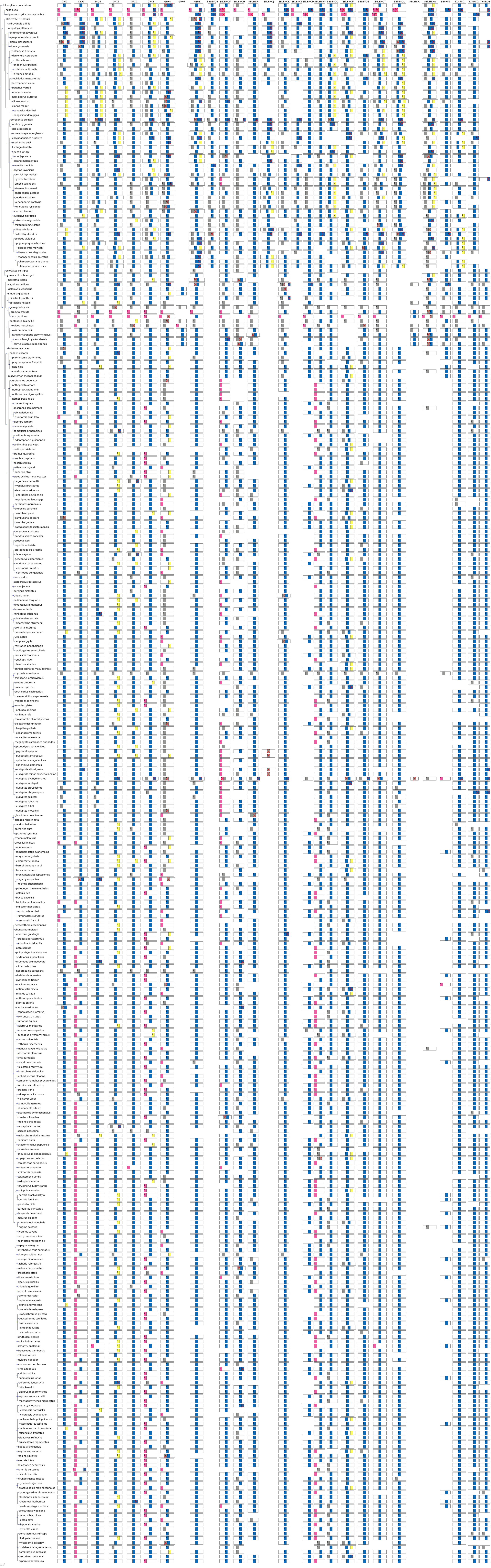
